# Identification of functional *Npu* DnaE and gp41-1 inteins split in three fragments

**DOI:** 10.1101/2023.01.30.526203

**Authors:** Daniel Weis, Navaneethan Palanisamy, Jara Ballestin Ballestin, Mehmet Ali Öztürk, Barbara Di Ventura

## Abstract

Inteins are special proteins that auto-catalytically carry out a protein splicing reaction. Due to their ability to post-translationally modify target proteins *in vitro* and *in vivo*, they are used in different applications, ranging from protein purification to the construction of Boolean logic gates. So far inteins have been found to be either encoded by a single gene (contiguous inteins) or by two separate ones (split inteins). Previously, it has been shown that the contiguous *Ssp* and *Rma* DnaB inteins and the split *Npu* DnaE intein could be artificially split in three fragments and retain functionality. Here we report the identification of novel split sites within the N-terminal fragments of the *Npu* DnaE and gp41-1 split inteins that lead to synthetic functional three-piece versions of these inteins. These variants contribute to the toolkit of three-piece inteins that could be used in biotechnological applications based on highly-fragmented inteins.

## Introduction

Inteins are proteins that autocatalytically excise themselves out of precursor host proteins cleaving the peptide bonds connecting them to the N- and C-terminal flanking sequences (exteins) and creating a new peptide bond to ligate these flanking sequences forming the splice product [1]. Inteins have been found in over six hundred organisms in all domains of life, and are usually embedded in genes encoding proteins essential for DNA replication and/or transcription, or cell homeostasis [2].

The inteins that have been so far identified in genomes are either encoded by a single gene or by two separate ones [3] [4]. The two fragments of split inteins (which we will refer to as N-intein and C-intein) spontaneously form a complex and perform a *trans*-splicing reaction following what has been, at least for the *Npu* DnaE intein, reported to be a “capture and collapse” mechanism [5] [6]. It is likely that split inteins evolved from a contiguous one after genomic rearrangement(s) that led to the separation of the coding sequence into two parts [4]. Similar to contiguous inteins, split inteins vary in sequence, but are highly similar in their overall structure and, interestingly, split site [7].

Contiguous and split inteins can be used to post-translationally modify target proteins *in vitro* and *in vivo* and are becoming more and more popular in the synthetic biology/protein engineering community. In particular, split inteins allow for a larger variety of applications compared with contiguous ones. They facilitate the addition or removal of peptide sequences from a protein of interest (POI) [8]; they can be employed to circularize POIs to increase their stability and decrease their aggregation propensity [9]–[11]; and they can be employed to construct AND gates in synthetic circuits [12].

If inteins split into two fragments exist, one can ask whether inteins split in more than two fragments also exist. While this has not been so far proved to occur in nature –likely due to the difficulty of identifying short DNA sequences that represent indeed intein fragments–, it has been experimentally shown to be possible: the contiguous inteins *Ssp* and *Rma* DnaB as well as the split intein *Npu* DnaE were successfully split into three fragments (N-, M-, and C-intein) [6] [13] [14]. Interestingly, the *Rma* DnaB intein split in three fragments was employed to construct a TALE (Transcriptional activator-like effectors)-based three-input AND circuit [14]. This 3 input-AND gate did show some activity when only the N and C-inteins were present (only 2 instead of 3 inputs), most likely due to the splicing being possible even in the absence of the M-intein [14]. To obtain more reliable 3-input intein-based AND gates, additional inteins that retain activity when split in three fragments, and for which no *trans*-splicing is observed in the absence of the M-intein, are required.

Here we show that the N-terminal fragment of the naturally split *Npu* DnaE (Npu) and gp41-1 inteins can be further split into two parts leading to three-piece inteins that retain functionality and strictly depend on the presence of all three fragments. These new highly-fragmented inteins pave the way for more complex applications as well as give more flexibility to control the splicing reaction in space and time.

## Results

To find potentially functional split sites in the N-terminal fragment of Npu and gp41-1 we followed a similar strategy as previously published [15] and looked for residues located within or near flexible loops and between β-sheets that could favour non-covalent bond formation without affecting the overall protein structure. We selected 7 split sites for gp41-1 (referred to as gp41-1 S1-S7; Figure 1 a, b) and 6 split sites for Npu (referred to as Npu S1-S6; Figure 1 c, d). Splitting the N-intein into two fragments leads to the formation of a new, shorter N-intein and a middle intein (M-intein). For *trans*-splicing to occur, three parts must come together: the N-, M- and C-inteins. To test the resulting three-piece inteins, we selected as model proteins to ligate together the maltose binding protein (MBP; N-extein) and thioredoxin (TRX; C-extein), since these two proteins are not expected to interact with each other contributing to the formation of the splicing-competent tri-molecular complex (Figure 2a).

**Figure 1.**
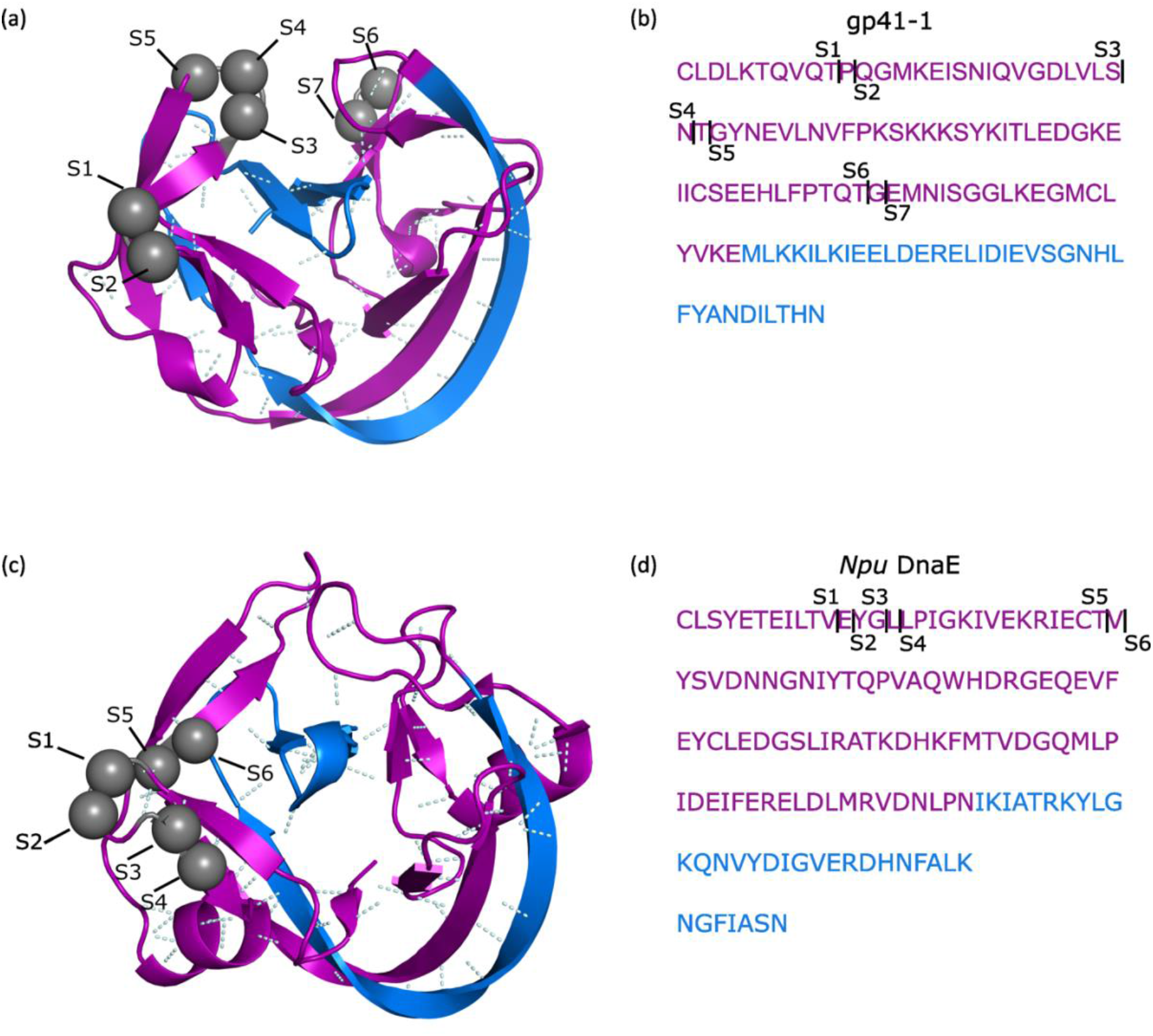
Location of new split sites in gp41-1 and *Npu* DnaE. (**a**,**c**) Crystal structure of gp41-1 (**a**; PDB id: 6QAZ) and *Npu* DnaE (**c**; PDB id: 4KL5) with the identified split sites shown as grey spheres. Non-covalent bonds are shown with pale cyan dashed lines. (**b**,**d**) Amino acid sequence of gp41-1 (**a**) and *Npu* DnaE (**d**). Split sites are indicated by black arrows and the letter S, with increasing subscript from left to right. Black numbers indicate amino acid positions. (**a**-**d**) The natural N- and C-inteins are shown in purple and blue, respectively.

**Figure 2.**
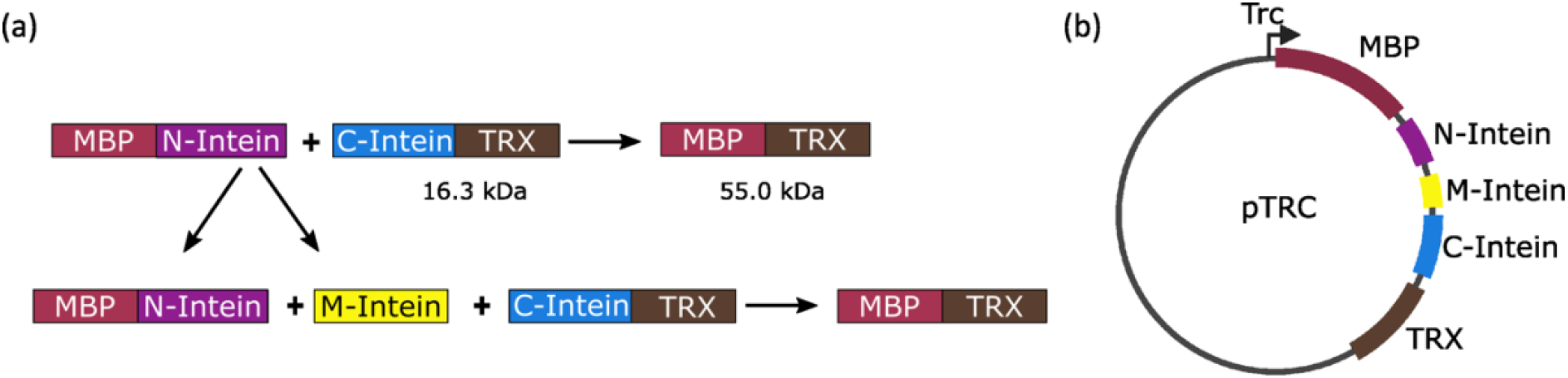
Construct used to test the three-piece inteins. (**a**) Schematic representation of the protein *trans*-splicing reaction for a two-(upper panel) and three-piece (lower panel) intein. MBP, maltose binding protein. TRX, thioredoxin. (**b**) Scheme of the plasmid used for testing different split sites. The multi-cistronic construct is driven by the pTrc promoter.

First, we cloned the genes encoding the fusion protein of MBP and the N-intein, and the fusion protein of the C-intein and TRX separated by a spacer containing a STOP codon, a ribosome binding site (RBS) and a start codon into a vector bearing an IPTG-inducible promoter (Figure 2b). We then cloned a similar spacer at the selected positions to create the split version of the N-intein (giving rise to the N- and M-inteins; Figure 2b). To assess the necessity to have all three parts for the splice reaction to occur, we cloned versions of these vectors lacking either the N-, M-or C-intein. Additionally, we cloned the positive controls MBP-Npu-TRX and MBP-gp41-1-TRX, where the naturally split inteins have been artificially made contiguous.

The plasmids were introduced into *E. coli* TOP10 cells and the expression of the constructs was induced with IPTG for 3 hours. The MBP-N-intein and C-intein-TRX constructs, as well as the MBP-TRX splice product were detected with antibodies via Western blotting. We decided not to add any tag to the M-intein, to introduce the least disturbance possible. As a consequence, the M-intein could not be detected, and statements about splicing efficiency could not be made.

All 7 split sites for gp41-1 allowed *trans*-splicing to occur (Figure 3a and Supplementary Figure S1). Interestingly, only Npu S1-S4 were associated with *trans*-splicing activity (Figure 3b and Supplementary Figure S2). We confirmed that all three pieces of the inteins are needed for the splicing to occur (Supplementary Figures S3 and S4).

**Figure 3.**
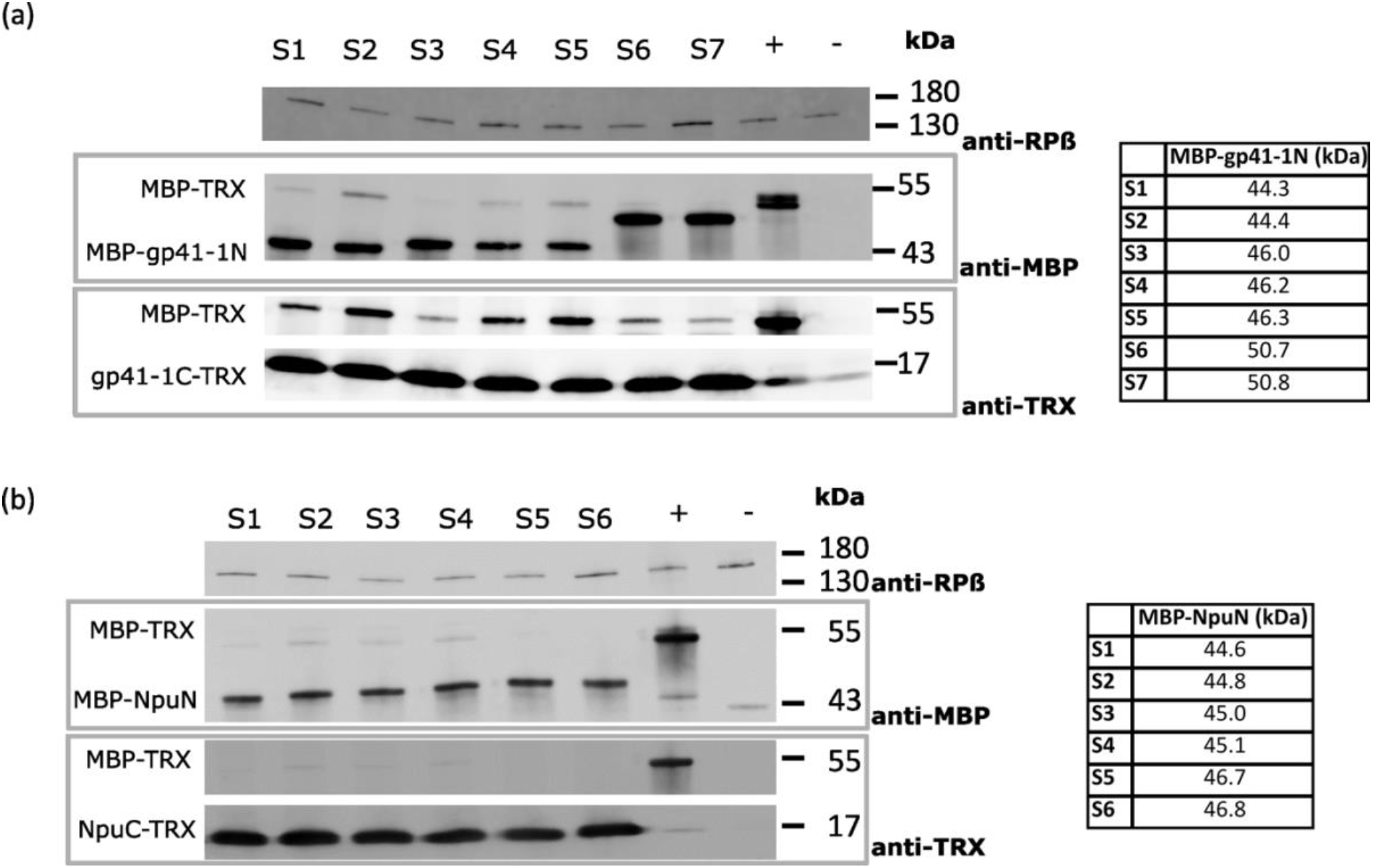
gp41-1 and *Npu* DnaE can be split in three fragments and retain *trans*-splicing activity. (**a**,**b**) Representative Western blot showing the formation of splice product for gp41-1 (**a**) and *Npu* DnaE (**b**) split at the indicated sites. +, positive control consisting of MBP-gp41-1-TRX (**a**) and MBP-Npu-TRX (**b**). -, negative control consisting of *E. coli* TOP10 cells transformed with empty pTrc99a. The experiment was performed three times with similar results. See Supplementary Figure 1 and 2 for full Western Blots.

## Discussion

In this study we showed that the *Npu* DnaE and gp41-1 inteins can be split into three fragments and still retain *trans*-splicing activity. The same was previously done with the artificially split *Ssp* and *Rma* DnaB inteins [14] [16] and the naturally split Npu DnaE intein [6]. Our results demonstrate that the ability to function when split in three pieces is not unique to these inteins, but can also be applied to the naturally split gp41-1 intein, which is the fastest split intein identified to date.

Our current data do not allow us to draw any conclusion about efficiency of splicing of the engineered three-piece inteins. For this, we would need to detect also the M-intein. At this stage, we cannot exclude that the weaker splice product observed compared with the positive control be due to very low M-intein levels. We decided to leave the M-intein untagged because we observed problems in another set of experiments (performed in mammalian cells) when fusing other proteins (such as GFP) to the M-intein of gp41-1 (unpublished results). Our data prove that gp41-1 and *Npu* DnaE –split at different sites compared to the one previously tested [6]– can be split in three pieces. Further studies are needed to assess the efficiency or boost it, were it too low for applications.

## Material and Methods

### Split site selection

We used the crystal structures of gp41-1 and *Npu* DnaE (PDB id: 6QAZ and PDB id: 4KL5, respectively) and identified loop regions and non-covalent bonds using PyMOL [17].

### Plasmid construction

To clone MBP (maltose binding protein) into pTrc99a, the *mbp* gene was amplified by PCR from plasmid pETM41 (including the TEV protease cleavage site) using a forward primer containing the NcoI restriction site and a reverse primer containing the BamHI restriction site. To clone TRX (thioredoxin) into the same pTrc99a, the *trxA* gene from plasmid pETTrx was amplified using a forward primer containing the BamHI restriction site and a reverse primer containing the PstI restriction site. Plasmids pETM41 and pETTrx were a kind gift of Gunter Stier (Heidelberg University Biochemistry Center). The backbone and the PCR fragments were digested with the corresponding restriction enzymes and then ligated together yielding plasmid pTrc-MBP-TRX. In addition to the above-mentioned restriction site, the reverse primer of TRX also carried an ochre STOP codon. Supplementary Table S1 lists the primers used in this study. Upon sequencing the pTrc-MBP-TRX construct, we found two instead of one TEV protease cleavage site between MBP and TRX. We nonetheless went on with this construct since we reasoned the second TEV cleavage site represented an additional linker and would not interfere with the conclusions of this study. The full-length gp41-1/*Npu* DnaE genes were cloned between the *mbp* and *trxA* genes in the pTrc-MBP-TRX construct via Gibson Assembly® yielding the pTrc-MBP-gp41-1-TRX and pTrc-MBP-Npu-TRX constructs. An ochre STOP codon and a ribosome binding site (RBS) were introduced between the N- and C-inteins in the pTrc-MBP-gp41-1-TRX and pTrc-MBP-Npu-TRX constructs performing a PCR with phosphorylated primers followed by blunt-end ligation. Finally, using the same blunt-end ligation method, split sites were added in the N-intein by introducing an amber STOP codon and an RBS.

### Bacterial growth and induction conditions

Plasmids were transformed into One Shot™ *E. coli* TOP10 cells from ThermoFisher. A single colony was picked and grown over night in LB medium at 37°C with shaking. The following morning the OD was measured and used to adjust the volume used for inoculation. The bacteria were then grown in LB medium for 1 hour and 30 minutes at 37°C with shaking before IPTG was added to a final concentration of 1 mM. OD was measured again to adjust the volume of samples taken. Subsequently, the bacteria were pelleted and resuspended in Laemmli buffer, and lysed by boiling for 10 minutes at 95°C. The samples were stored at -20°C.

### Western Blot

Samples were loaded on a 10% SDS-PAGE gel and run at 100 V for 75 minutes. Proteins were then transferred onto a PMSF membrane using semi-dry blotting. The membrane was blocked with BSA and then incubated with the primary antibodies (see **Table 1**) overnight at 4°C with gentle rocking. The membrane was then washed in TBS-Tween and incubated with the secondary antibodies (see **Table 2**) for 1 hour at room temperature with gentle rocking. After further washing with TBS-Tween, the membrane was imaged in an Amersham Typhoon laser scanner.

**Table 1:** List of primary antibodies

| Antibody | Species | Dilution | Manufacturer (Product ID) |
| --- | --- | --- | --- |
| Anti-MBP | mouse | 1:10000 | NEB (E8032L) |
| Anti-TRX | rabbit | 1:10000 | Invitrogen (PA5-117517) |
| Anti-RNA polymerase $\beta$ | mouse | 1:1000 | Biolegend (662904) |

**Table 2.** List of secondary antibodies

| Antibody | Dye | Dilution | Manufacturer (Product ID) |
| --- | --- | --- | --- |
| Anti-mouse | Alexa Fluor™ 488 | 1:2000 | Invitrogen (A-11029) |
| Anti-rabbit | Cy5 | 1:2000 | Invitrogen (A-10523) |

## Acknowledgments

This study was funded by the DFG (grant no. VE776/3-2 to B.D.V. within the SPP1926) and by the Excellence Initiatives of the German Federal and State Governments BIOSS (Centre for Biological Signalling Studies; EXC-294), and CIBSS (Centre for Integrative Biological Signalling Studies; EXC-2189).

## Author contribution

D. W. conceived the idea. N.P. cloned the constructs. D.W. and J.B. performed experiments. M.A.Ö. performed split site prediction. B.D.V. provided funding, supervised the study and wrote the manuscript. All authors discussed the data and approved of the manuscript.

