## Supplementary Information for "Identification of functional *Npu* DnaE and gp41-1 inteins split in three fragments"

for

### Table of content

Supplementary Figures S1-S4

Supplementary Table S1

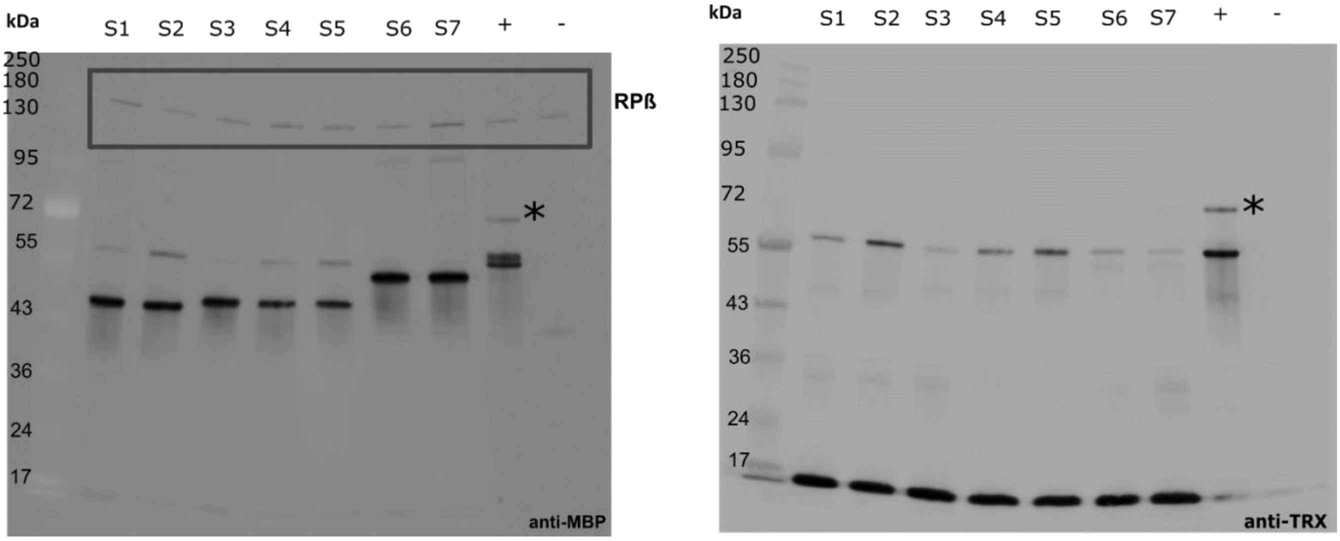


**Supplementary Figure S1**. Full Western Blot for Figure 3a. Asterisks indicate the precursor fusion protein MBP-gp41-1-TRX, still carrying the intein (molecular weight of 69 kDa). This indicates that the contiguous version of gp41-1 does not fully splice itself out of the precursor protein. RPβ, RNA polymerase beta subunit, used as loading control.

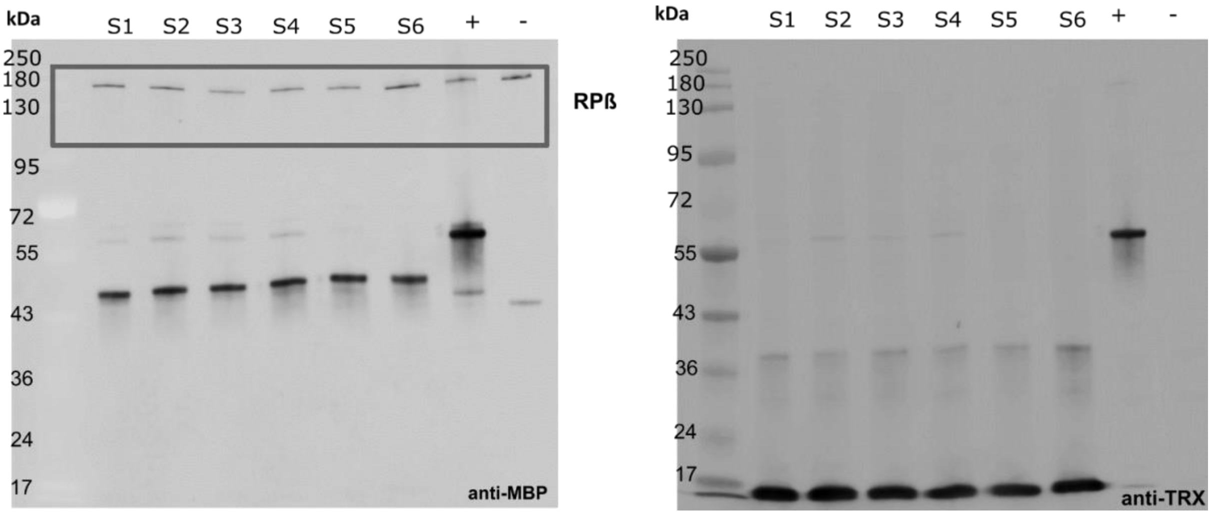


**Supplementary Figure S2**. Full Western Blot for Figure 3b. RPβ, RNA polymerase beta subunit, used as loading control.

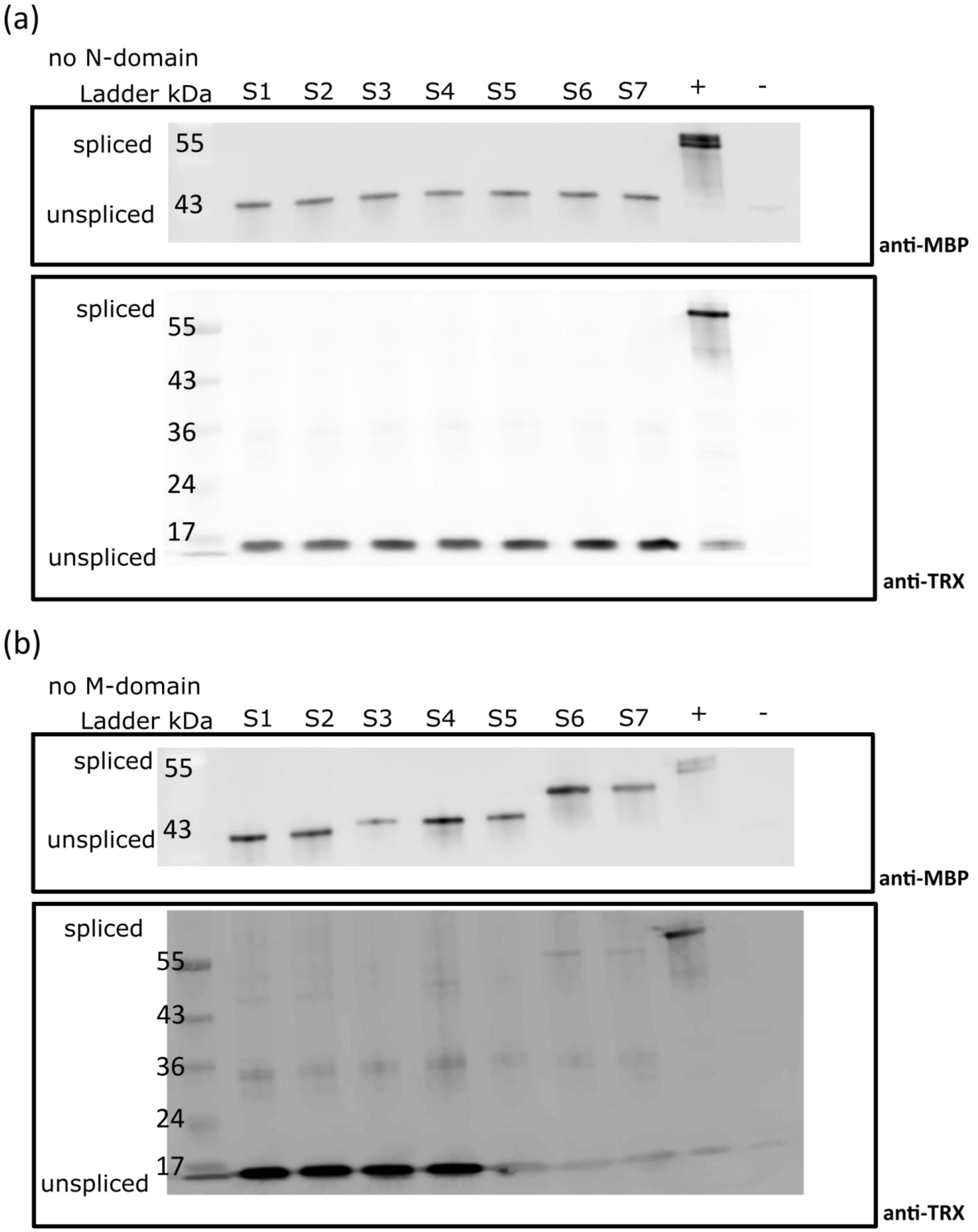
(a)
no N-domain
Ladder kDa
S1
S2
S3
S4
S5
S6
S7
+
-
spliced
unspliced
55
43
anti-MBP
spliced
unspliced
55
43
36
24
17
anti-TRX
(b)
no M-domain
Ladder kDa
S1
S2
S3
S4
S5
S6
S7
+
-
spliced
unspliced
55
43
anti-MBP
spliced
unspliced
55
43
36
24
17
anti-TRX

(c)

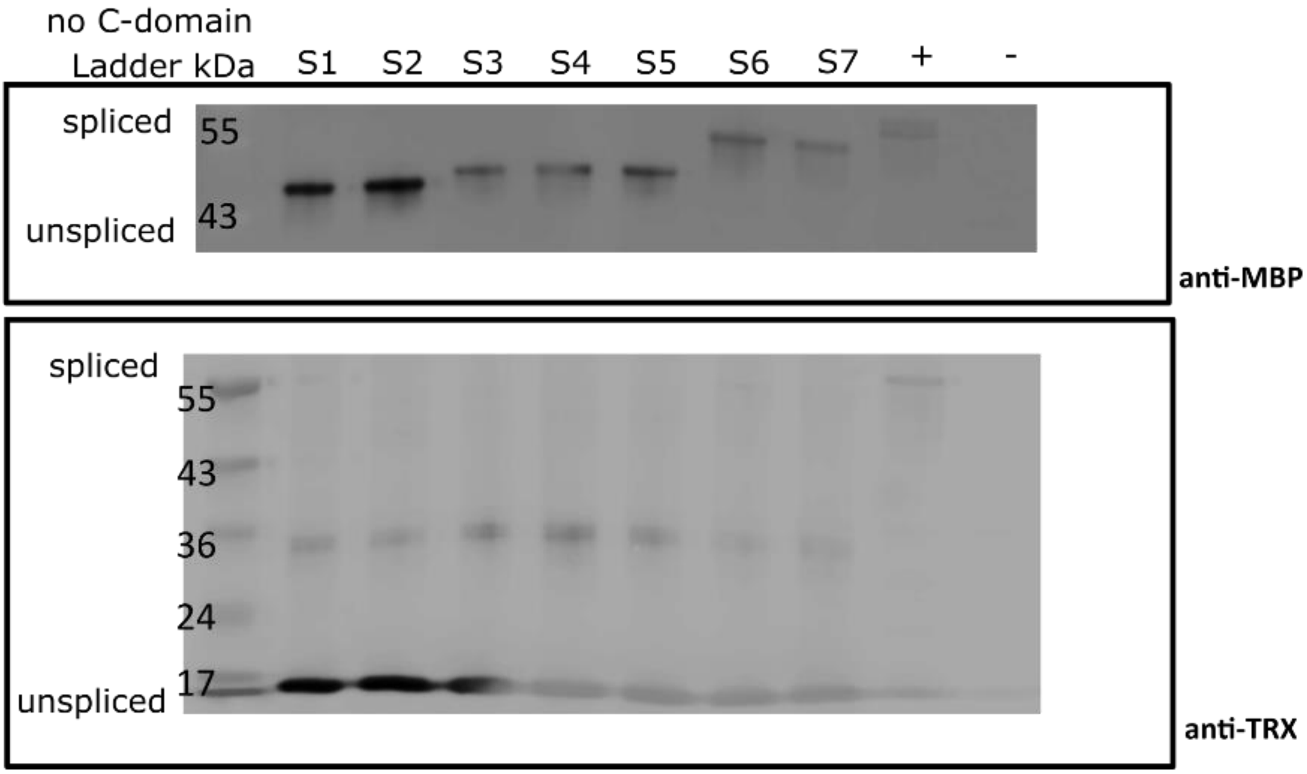


**Supplementary Figure S3**. All three gp41-1 pieces are necessary for the splicing reaction. (**a**-**c**) Representative Western Blot showing the splice product (spliced) and the precursors (unspliced) when either the N- (**a**), the M- (**b**) or the C-intein (**c**) are not expressed. +, positive control consisting of *E. coli* TOP10 cells transformed with the MBP-gp41-1-TRX construct. -, negative control consisting of *E. coli* TOP10 cells transformed with an empty pTrc99a plasmid.

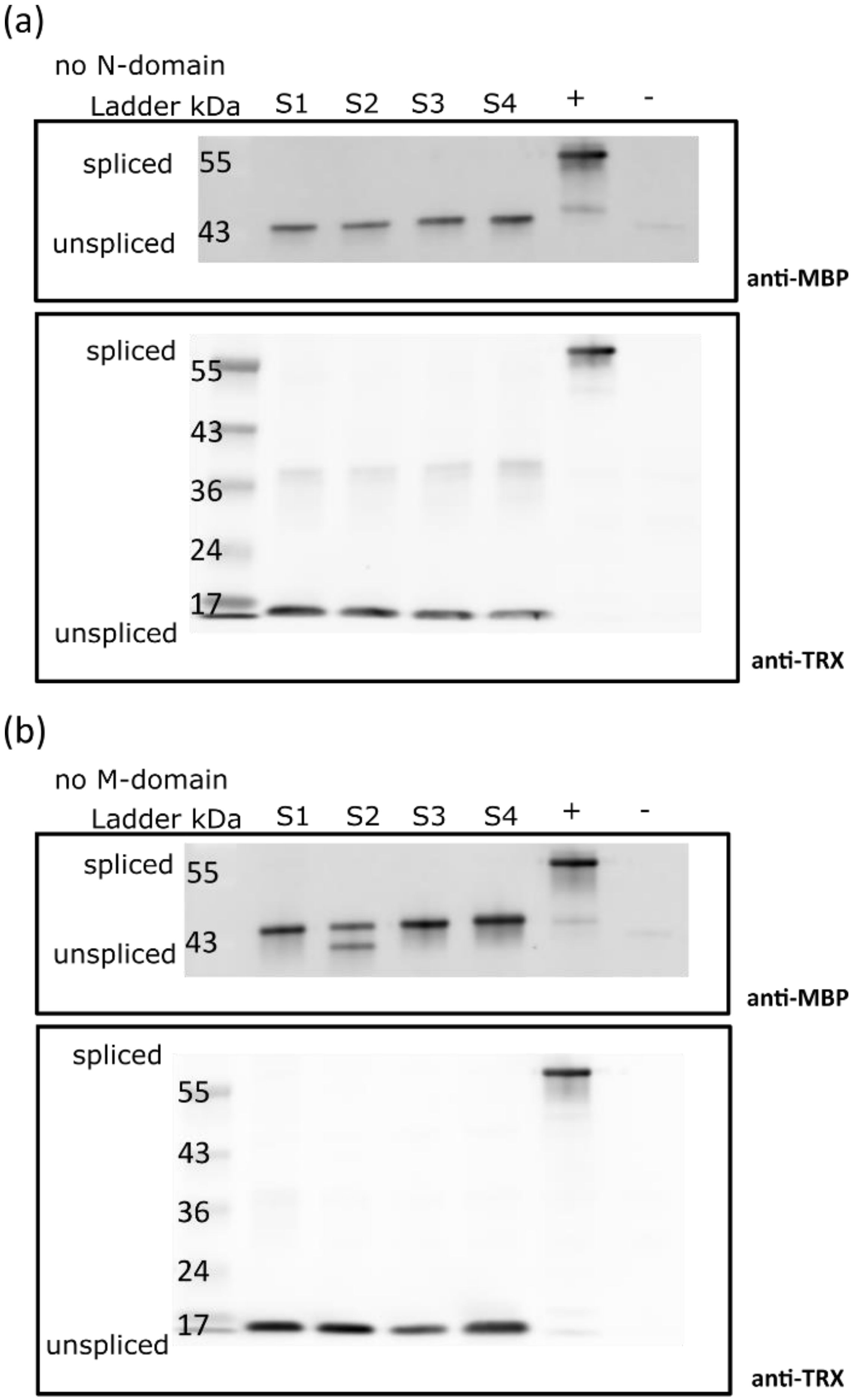
(a)
no N-domain
Ladder kDa
S1
S2
S3
S4
+
-
spliced
55
unspliced
43
anti-MBP
spliced
55
43
36
24
17
unspliced
anti-TRX
(b)
no M-domain
Ladder kDa
S1
S2
S3
S4
+
-
spliced
55
unspliced
43
anti-MBP
spliced
55
43
36
24
17
unspliced
anti-TRX

(c)

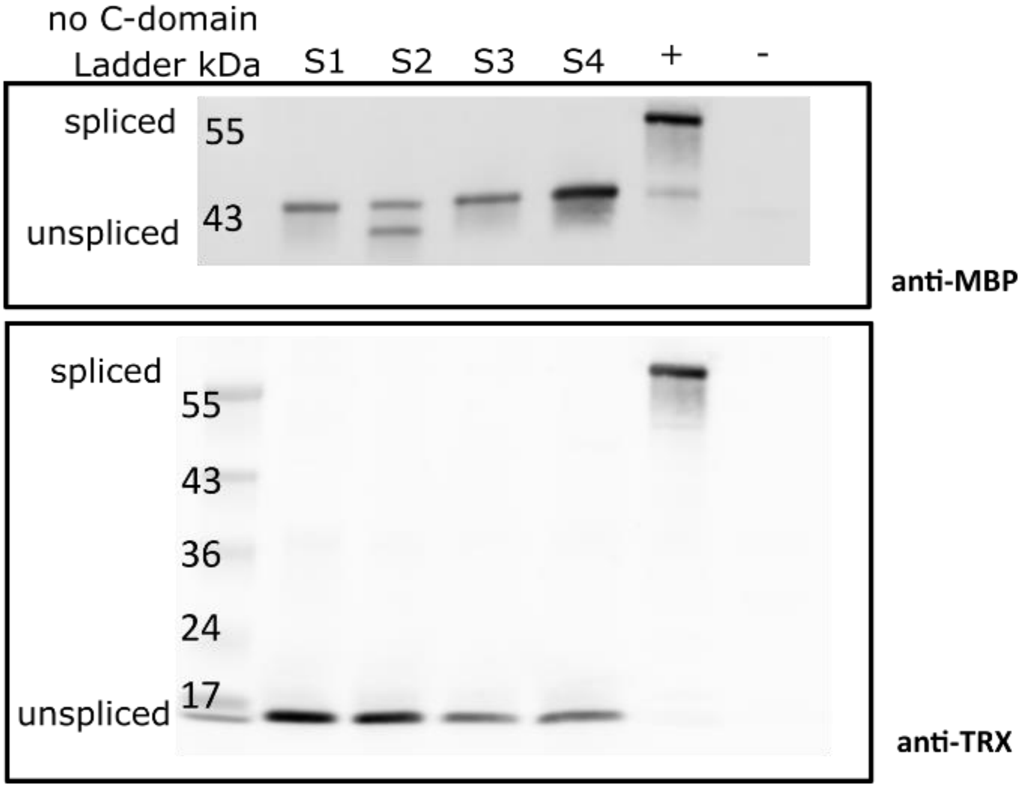


**Supplementary Figure S4**. All three *Npu* DnaE pieces are necessary for the splicing reaction. (**a-c**) Representative Western Blot showing the splice product (spliced) and the precursors (unspliced) when either the N- (**a**), the M- (**b**) or the C-intein (**c**) are not expressed. +, positive control consisting of *E. coli* TOP10 cells transformed with the MBP-Npu-TRX construct. -, negative control consisting of *E. coli* TOP10 cells transformed with an empty pTrc99a plasmid.

**Supplementary Table S1**. List of primers used in the present study

| Primer name | Sequence (5’ to 3’) |
|---|---|
| NcoI_MBP_FP | ttt*ccatgg*aaatcgaagaaggtaaactggtaatctggattaacggcg |
| BamHI_TEV_MBP_RP | aaa*ggatcc*gccctgaaaataaagattctcgctca |
| BamHI_TRX_FP | aaa*ggatcc*atgagcgataaaattattcacctgactgacgacagttttgacacgg |
| PstI_TAA_TRX_RP | aaa*ctgcag*ttaggccaggttagcgtcgaggaactctttcaactgacc |
| Gib_MBP-TRX_FP | atgagcgataaaattattcacctgactgacgacagttttgacacgg |
| Gib_MBP-TRX_RP | gccctgaaaataaagattctcgctcatgg |
| Gib_gp41_N_FP | gagcgagaatctttattttcagggctgcttggatctgaaaacccaggttcagaccccgc |
| Gib_gp41_N_RP | ttctttaacatacagacacataccttctttcagaccaccgg |
| Gib_gp41_C_FP | aggtatgtgtctgtatgttaaagaaatgctgaaaaaaatcctgaaaatcgaagagctgg |
| Gib_gp41_C_RP | gaataattttatcgctcatggatccagagttgtgggtcagaatatcgttagcgtaaaacagg |
| Gib_npu_N_FP | gagcgagaatctttattttcagggcaggggcaagtgcttgtcctacgaaaccg |
| Gib_npu_N_RP | ggtacttgcgggtggcgatcttgatgttgggcaggttgtccacgcgc |
| Gib_npu_C_FP | atcaagatcgccacccgcaagtacctgg |
| Gib_npu_C_RP | gaataattttatcgctcatggatccctcccagcagttggaggcgatgaagc |
| Phos_gp41_RBS_FP | aataattttgtttaactttaagaaggagatataccatgctgaaaaaaatcctgaaaatcgaagagctgg |
| Phos_gp41_TAA_RP | ttattctttaacatacagacacataccttctttcagaccaccgg |
| Phos_npu_TAA_RBS_FP | taaaataattttgtttaactttaagaaggagatataccatgatcaagatcgccacccgcaagtacc |
| Phos_npu_RP | gttgggcaggttgtccacgcgcatcagg |
| Phos_gp41_11_FP | tagataaaggaggtaaataatgcagggtatgaaggaaatttccaacatccaggtcggtgatctgg |
| Phos_gp41_11_RP | cggggtctgaacctgggttttcagatccaagca |
| Phos_gp41_12_FP | tagataaaggaggtaaataatgggtatgaaggaaatttccaacatccaggtcggtgatctggtactg |
| Phos_gp41_12_RP | ctgcggggtctgaacctgggttttcagatccaagca |
| Phos_gp41_28_FP | tagataaaggaggtaaataatgaacacgggttacaacgaagttctgaacgtcttcccg |
| Phos_gp41_28_RP | gctcagtaccagatcaccgacctggatgttgg |

| | |
|---|---|
| Phos_gp41_29_FP | tagataaaggaggtaaataatgacgggttacaacgaagttctgaacgtcttcccgaaatc |
| Phos_gp41_29_RP | gttgctcagtaccagatcaccgacctggatgttgg |
| Phos_gp41_30_FP | tagataaaggaggtaaataatgggttacaacgaagttctgaacgtcttcccgaaatctaaaaaaaag |
| Phos_gp41_30_RP | cgtgttgctcagtaccagatcaccgacctggatgttgg |
| Phos_gp41_69_FP | tagataaaggaggtaaataatgggtgaaatgaacatctccggtggtctgaaagaaggtatg |
| Phos_gp41_69_RP | agtctgcgtcggaaacaggtgttcttcggaacag |
| Phos_gp41_70_FP | tagataaaggaggtaaataatggaaatgaacatctccggtggtctgaaagaaggtatgtgtctg |
| Phos_gp41_70_RP | accagtctgcgtcggaaacaggtgttcttcggaacag |
| Phos_npu_11_FP | tagataaaggaggtaaataatggagtacggcctgctgcccatcggc |
| Phos_npu_11_RP | cacggtcaggatctcggtttcgtaggacaagca |
| Phos_npu_12_FP | tagataaaggaggtaaataatgtacggcctgctgcccatcggcaag |
| Phos_npu_12_RP | ctccacggtcaggatctcggtttcgtaggacaagca |
| Phos_npu_14_FP | tagataaaggaggtaaataatgctgctgcccatcggcaagatcgtggagaagc |
| Phos_npu_14_RP | gccgtactccacggtcaggatctcggtttcgtaggacaagca |
| Phos_npu_15_FP | tagataaaggaggtaaataatgctgcccatcggcaagatcgtggagaagcgc |
| Phos_npu_15_RP | caggccgtactccacggtcaggatctcggtttcgtaggacaagca |
| Phos_npu_29_FP | tagataaaggaggtaaataatggtgtactccgtggacaacaacggcaacatctacaccc |
| Phos_npu_29_RP | ggtgcactcgatgcgcttctccacgatc |
| Phos_npu_30_FP | tagataaaggaggtaaataatgtactccgtggacaacaacggcaacatctacacccagcccg |
| Phos_npu_30_RP | cacggtgcactcgatgcgcttctccacgatc |
| Phos_delete_N_intein_FP | tagataaaggaggtaaataatg |
| Phos_delete_N_intein_RP | cgaattagtctgcgc |
| Phos_delete_M_intein_FP | cattatttacctcctttatcta |
| Phos_delete_M_intein_RP | taaaataattttgtttaactttaaga |
| Phos_delete_C_intein_FP | atgagcgataaaattattcacctgactg |
| Phos_delete_C_intein_RP | catggtatatctccttcttaaagttaaacaaaattatt |

Restriction site is represented in *italics*
